# Software Package for Transcranial Magnetic Stimulation Coil and Coil Array Analysis and Design

**DOI:** 10.1101/2023.08.20.554037

**Authors:** Leah Morales, William A Wartman, Jonathan Ferreira, Alton Miles, Mohammad Daneshzand, Hanbing Lu, Aapo R. Nummenmaa, Zhi-De Deng, Sergey N. Makaroff

**Affiliations:** Electrical and Computer Engineering, Worcester Polytechnic Inst., Worcester, MA 01609 USA; Analog Devices, Inc., 1 Analog Way, Wilmington, MA 01887 USA; A. A. Martinos Ctr., Massachusetts General Hospital, Harvard Medical School, Charlestown, MA 02129 USA; National Institute of Drug Abuse, NIH, Biomedical Research Center, 251 Bayview Boulevard, Baltimore, MD 21224 USA; Computational Neurostimulation Research Program, Noninvasive Neuromodulation Unit, Experimental Therapeutics & Pathophysiology Branch, National Institute of Mental Health, NIH, Bethesda, MD 20892-9663 USA

**Keywords:** Transcranial Magnetic Stimulation (TMS), TMS Coil Modeling and Design, TMS Coil Array Modeling and Design, Fast Multipole Method, MATLAB® platform

## Abstract

**Objective:** This study aims to describe a MATLAB software package for transcranial magnetic stimulation (TMS) coil analysis and design.

**Approach:** Electric and magnetic fields of the coils as well as their self- and mutual (for coil arrays) inductances are computed, with or without a magnetic core. Solid and stranded (Litz wire) conductors are also taken into consideration. The starting point is the centerline of a coil conductor(s), which is a 3D curve defined by the user. Then, a wire mesh and a computer aided design (CAD) mesh for the volume conductor of a given cross-section (circular, elliptical, or rectangular) are automatically generated. Self- and mutual inductances of the coil(s) are computed. Given the conductor current and its time derivative, electric and magnetic fields of the coil(s) are determined anywhere in space.

Computations are performed with the fast multipole method (FMM), which is the most efficient way to evaluate the fields of many elementary current elements (current dipoles) comprising the current carrying conductor at a large number of observation points. This is the major underlying mathematical operation behind both inductance and field calculations.

**Main Results:** The wire-based approach enables precise replication of even the most complex physical conductor geometries, while the FMM acceleration quickly evaluates large quantities of elementary current filaments. Agreement to within 0.74% was obtained between the inductances computed by the FMM method and ANSYS Maxwell 3D for the same coil model. Although not provided in this study, it is possible to evaluate non-linear magnetic cores in addition to the linear core exemplified. An experimental comparison was carried out against a physical MagVenture C-B60 coil; the measured and simulated inductances differed by only 1.25%, and nearly perfect correlation was found between the measured and computed E-field values at each observation point.

**Significance:** The developed software package is applicable to any quasistatic inductor design, not necessarily to the TMS coils only.

## I. INTRODUCTION

Transcranial magnetic stimulation (TMS) is a noninvasive neuromodulation technique that uses magnetic pulses to induce an electric field in the brain to stimulate specific cortical regions. TMS coil design is an important aspect of this technique, as the coil’s geometry and material properties determine the stimulation intensity, focality, and efficiency. TMS coil design can be performed with numerous finite element method (FEM) packages intended for quasistatic eddy current problems. For example, [1] used the FEM package MagNet (Infolytica, Inc., Canada) for a focality study of 50 different TMS coil designs. ANSYS Maxwell 3D [2],[3] or SEMCAD X [4] can be used as well. However, commercial FEM packages may be costly and are not without some limitations. For example, while ANSYS Maxwell 3D perfectly predicts the magnetic field, it might generate large errors for the induced electric field in air. Therefore, an accessible dedicated TMS coil design software may be useful.

Reference [5] developed a model of the TMS coil in the form of an ensemble of magnetic dipoles, which is used in the popular SimNIBS software [6],[7]. Recently, a collection of 25 such coil models has been recently made available with validated field measurements [8]. This model divides the area of a loop of current into subareas and places magnetic dipoles perpendicular to the loop area at the centers of the subareas. The dipoles are weighted by the loop current and by the subarea sizes.

A somewhat more natural model of the coil conductor is an ensemble of many elementary current filaments (current dipoles), comprising the current-carrying conductor. Both primary magnetic and electric fields of the coil can be computed from the Biot-Savart law and from the known magnetic vector potential. Furthermore, static inductance(s) can be straightforwardly computed using the well-known Neumann formula [9],[10],[11]. Wire-based coil models of specific coil geometries and of different complexities have been used for numerical optimization and experimental verification in TMS-related studies [12],[13],[14],[15],[16],[17],[18],[19],[20].

The goal of this study was to compile a useful and accessible MATLAB^®^ software package for TMS coil analysis and design. The package can compute electric and magnetic fields of the coils as well as their self- and mutual (for coil arrays) inductances, with or without a magnetic core. We use the wire-based approach for computations, which allows us to follow the physical conductor geometry and electric current distribution precisely. Both solid and stranded (Litz) wires can be modeled. Computations are performed with the fast multipole method or FMM (see [21],[22]) which is the most efficient way to evaluate the fields of many elementary current filaments (current dipoles) comprising the current-carrying conductor at a large number of observation points. This is the major underlying mathematical operation behind both inductance and field calculations; its speed up enables the organization of some large optimization loops.

As TMS continues to grow as a research tool and potential therapeutic intervention for various clinical indication, the development of efficient and accessible software for coil design will be increasingly important, and this work represents a valuable step forward in this area.

## II. Materials and Methods

### A. Software Workflow

Fig. 1 shows the coil design workflow. The design process begins with the user specifying a centerline of a coil conductor, which is a curve in three-dimensional space represented by an *N* × 3 array of nodes. If an array of coils is being designed, multiple curves can be given.

**Fig. 1.**
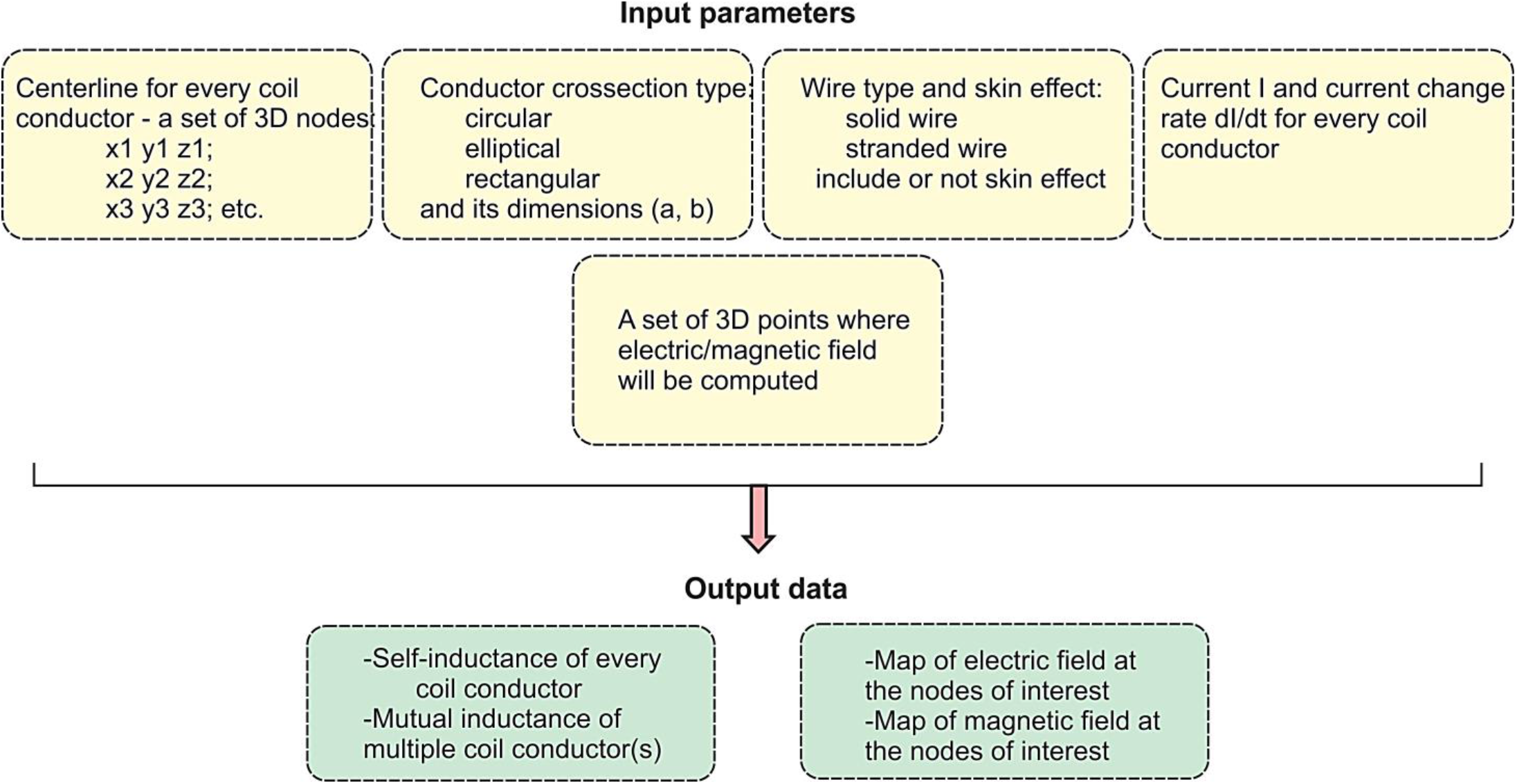
Design workflow of the coil (system) including input and output parameters.

The next step is to provide information about the conductor’s cross-section, including its type (circular, elliptical, or rectangular) and dimensions. Sweeping a non-symmetric cross-section along a curved centerline can be done in several ways. Therefore, a preferred orientation about the centerline’s directional vector should also be specified when a non-circular cross-section is used.

Given this information, an computational wire mesh and an illustrative computer aided design (CAD) mesh for the volume conductor will automatically be generated, as described in Section II.B. The wire mesh can model either solid or stranded (Litz wire) conductors. In the first case, the electric current will flow in the vicinity of the surface only (the skin effect). In the second case, it will be uniformly distributed throughout the conductor’s cross-section.

The electric and magnetic fields of the coil(s) can be determined anywhere in space by providing the conductor’s current and its time derivative, as described in Sections II.C and II.D, respectively. The fast multipole method is used to compute the fields at many observation points, which could be beneficial for optimization loops. Further, self- and mutual inductances of the coil(s) are also computed with the fast multipole method, as described in Section II.E.

Modifications necessary for the inclusion of a magnetic core are described in Section II.F. For the linear case, only the surface mesh of the core and its magnetic permeability are required. In the nonlinear case, an extra volumetric tetrahedral mesh for the core and a BH curve will be necessary. For simple core shapes, such meshes can be generated directly in MATLAB.

### B. Automated Generation of Computational Wire Coil Mesh and CAD Coil Mesh

There are various ways to sweep a cross-section along a path [23]. We create a triangular surface mesh using the method of a moving cross-section for the given centerline. Its concept is shown in Fig. 2a; it is applicable to any cross-sectional geometry with the primary nodes along its contour. Fig. 2a shows three points (labeled 1, 2, and 3) along the coil’s centerline. Two unit vectors are computed: **n**_1_ points from point 1 to point 2, and **n**_2_ points from point 2 to point 3. At point 2 we define a plane that passes through the point and whose unit normal vector **n**_**3**_ is given by the average of **n**_1_ and **n**_2;_ that is 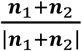.

**Fig. 2.**
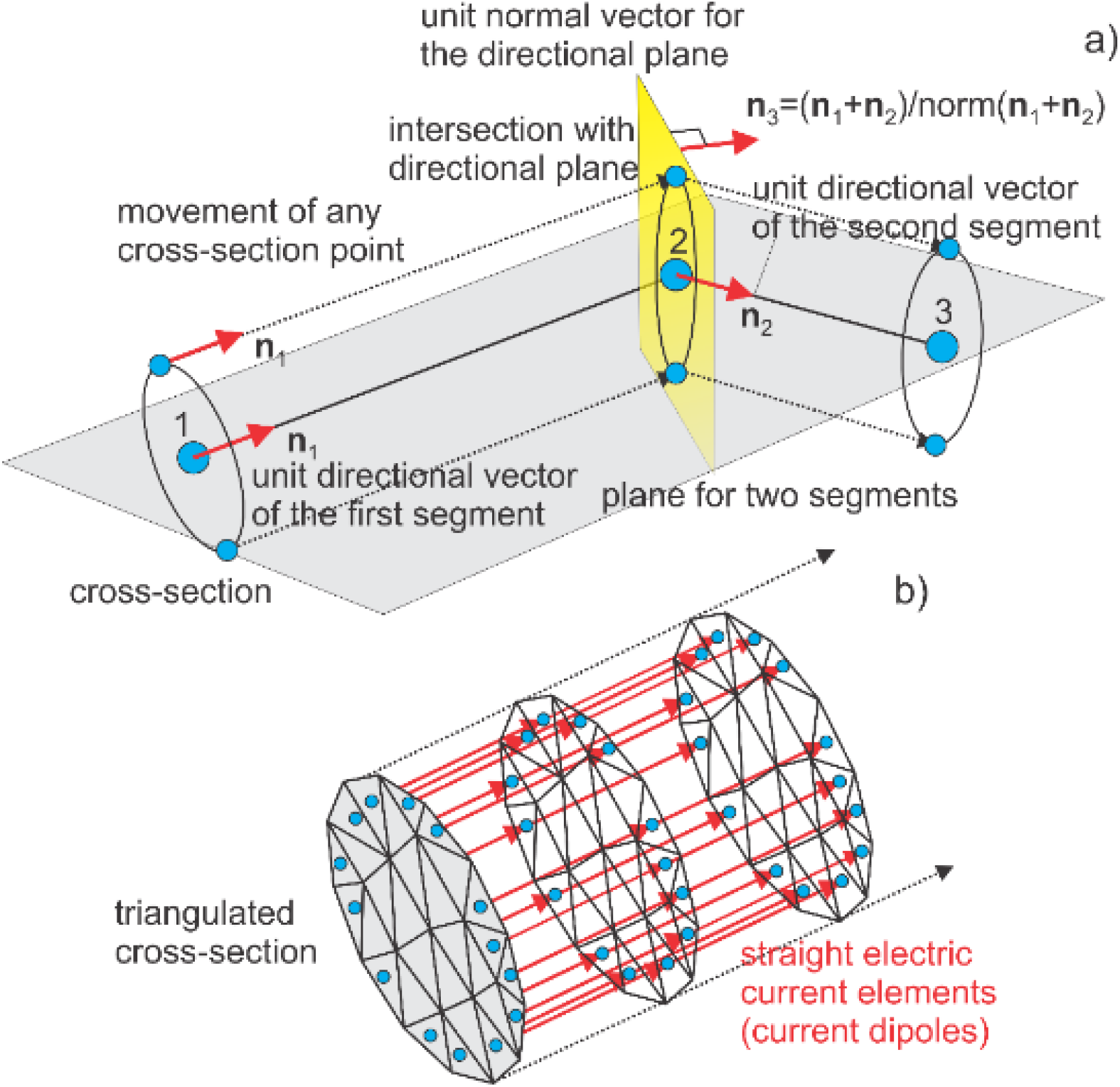
a) Method of a moving cross-section (or extrusion along a path) to create both wire and CAD meshes. The key is a directional plane, shown in yellow, that is formed for every two adjacent segments. This plane is used to extrude the cross-section. b) Formation of wire mesh by adding straight directional segments (elementary current dipoles), shown in red, at every extrusion step. For the case shown, these segments are only added at the boundary, which corresponds to a solid conductor with the skin effect.

The nodes of cross-section 1 in Fig. 2a are moved in the direction of the unit vector of segment 1 until they cross the directional plane found for point 2. The translated points form the new cross section about point 2. Triangles are then drawn to connect the outermost vertices of cross-sections 1 and 2, forming a triangular surface mesh. This process repeats for each centerline vertex, and the nodes and facets of every cross-section are accumulated. At the end, planar caps are introduced to obtain a 2-manifold CAD mesh.

The corresponding computational wire mesh is obtained by connecting centers of the facets of the sequential cross-sections as shown in Fig. 2b, thus creating small elementary current filaments (current dipoles). Either the border facets (the skin effect) or all facets (Litz wire) can be connected. Fig. 3, presented by [24], is an illustration of one combined (CAD + wire) coil mesh obtained in this way. The current filaments are always parallel to the centerline of the conductor. The coil conductor may be either closed or open as shown in Fig. 3. No bifurcations are supported at this point, but the bifurcation algorithm and software do exist and could be incorporated.

**Fig. 3.**
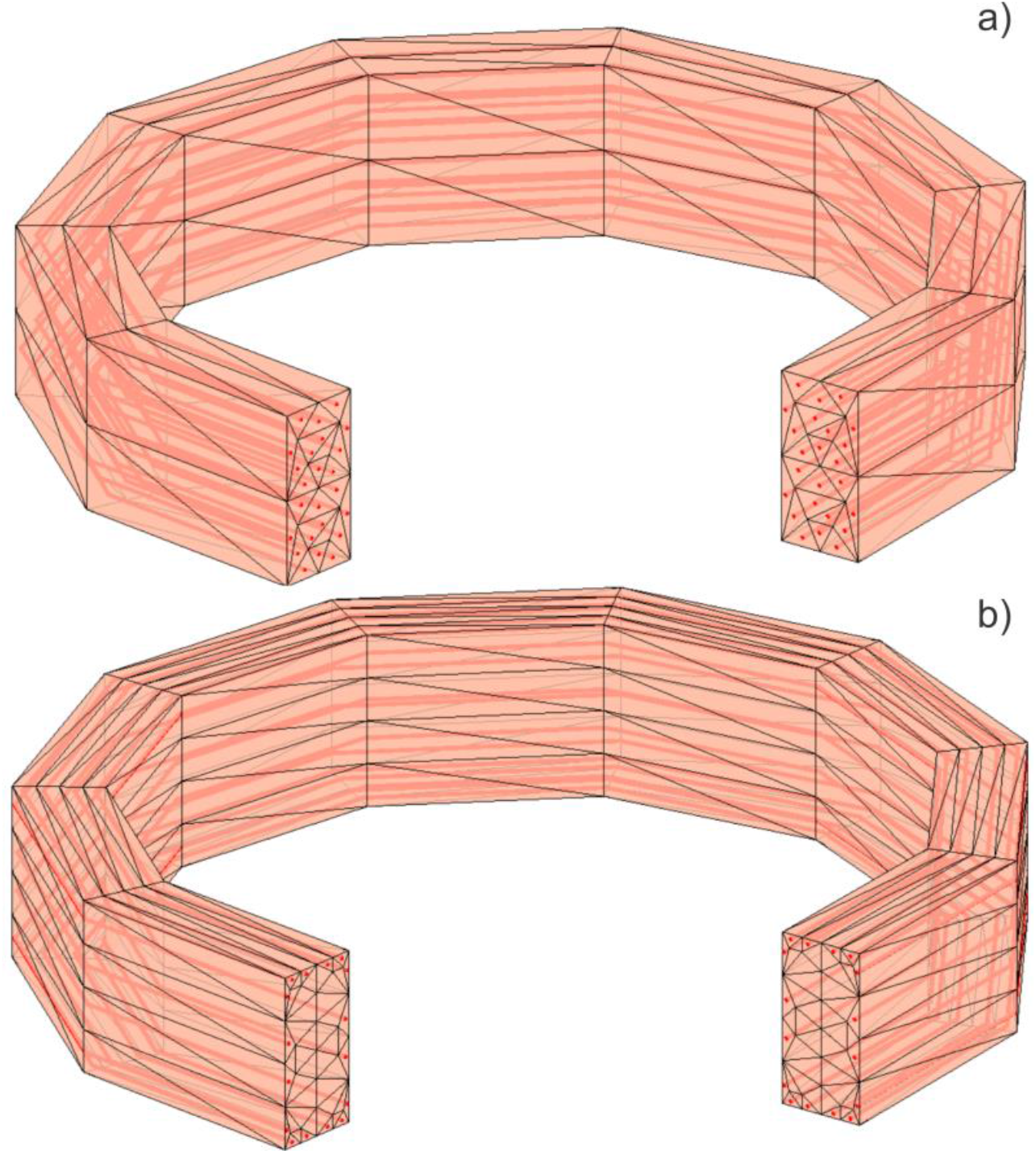
Combined (CAD + wire) coil mesh. Elementary wire segments used for actual computations are shown in red while edges of the surface facets are shown in black. a) Uniform current flow (Litz wire). b) Current flow in a solid conductor with the skin effect. The CAD mesh mainly serves illustration purposes.

For a non-circular cross-section and a non-planar centerline path, this method may be corrected to follow a desired cross-sectional orientation over the entire path. This is done by rotating the cross-sectional mesh in every directional plane so that the cross-section is always best aligned with a certain predefined direction, e.g., with the vertical direction. Simultaneously, additional user-defined rotations of the cross-section may be introduced at every sweeping step to describe, for example, a heavily twisted Litz wire.

### C. Induced Electric Field of the Coil

Consider a conductor that carries a total current *I*(*t*). Next, consider a small straight filament of current *I*_*n*_(*t*) = *I*(*t*)*i*_*n*_ within that conductor, which has vector length ***s***_*n*_ and is centered at ***r***_*n*_. Dimensionless scalar *i*_*n*_ < 1 is the “weight” of the filament; it is defined based on the wire mesh. If *S* filaments uniformly distributed in space were to pass through a conductor’s cross section, this weight could be equal to *1/S* for each of them.

The magnetic vector potential, ***A***, of this current filament at an observation point ***r***_*m*_ is given by [25]

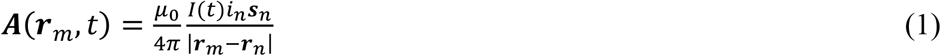

where *μ*_0_ is magnetic permeability of vacuum. Therefore, the electric field generated by this current filament is

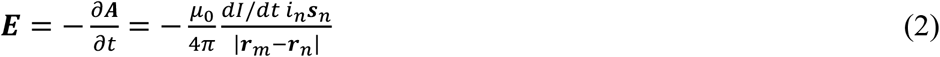

Now consider N current filaments or dipoles with dipole moments *i*_*n*_***s***_*n*_ acting on M observation points ***r***_*m*_. The electric field at any ***r***_*m*_ is given by

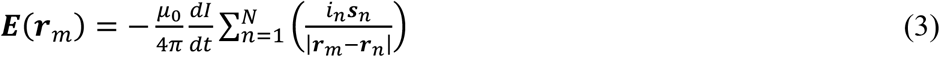

Expanding the quantity within the sum into its three Cartesian components,

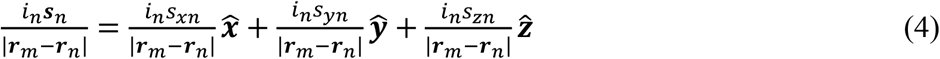

Each component has the form of the potential of a pseudo-electric charge *q*_(*x*/*y*/*z*)*n*_ = *i*_*n*_*S*_(*x*/*y*/*z*)*n*_, so the sum in Eq. (3) can be evaluated directly by applying the FMM to the three sets of pseudo charges.

### D. Magnetic Flux of the Coil

The same small straight filament of current generates a magnetic flux ***B*** given by Biot-Savart law [25]

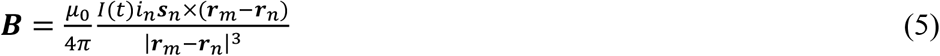

For multiple segments and multiple observation points, Eq. (5) cannot be evaluated using the FMM framework directly. However, it can be rewritten in the form

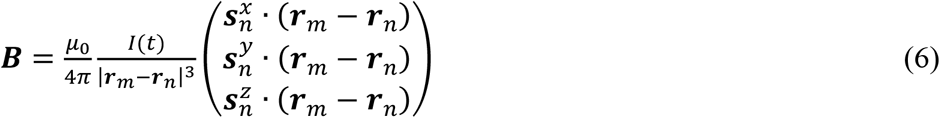

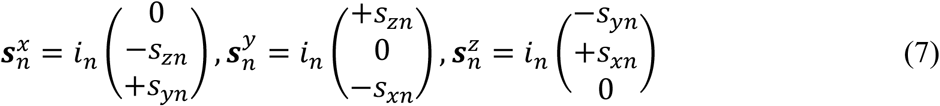

that allows one to use the standard FMM formulation for the electric potential of a double layer (layer of electric dipoles) three times. The dipole strengths are 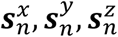. The FMM result is then multiplicatively scaled by *μ*_0_*I*(*t*)/(4*π*) as required by Eq. (6).

### E. Coil Inductance(s)

The magnetic energy U of an arbitrary current flow is expressed through the volumetric non-discretized current density ***j***(***r***) in the form [26]

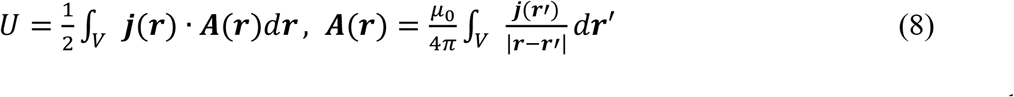

The (static) self-inductance of the coil, *L*, is found directly from the energy relation, 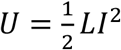, after substitution of Eqs. (8). Here, *I* is the total terminal (DC) coil current. If the volumetric current flow is discretized into *N* short straight current filaments with vector length ***s***_*n*_, each carrying current *i*_*n*_ and centered at ***r***_*n*_, the coil inductance without the core is given by the Neumann formula [9],[10],[11]

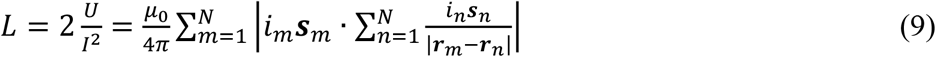

The inner sum in Eq. (9) is computed via the FMM as a potential of a single layer repeated three times – see Eqs. (3) and (4). Those computations are done in parallel. After that, the outer sum is computed directly. We found that for precise inductance calculations via the Neumann formula, the ratio of average segment length to the average segment spacing should be on the order of 1. The self-terms of the Neumann formula are ignored (set to zero).

For the mutual inductance *M* between the two coils, Eq. (9) still holds, but the factors *i*_*m*_***s***_*m*_ and *i*_*n*_***s***_*n*_ in Eq. (9) now relate to the two different coils with *M* and *N* filaments, respectively,

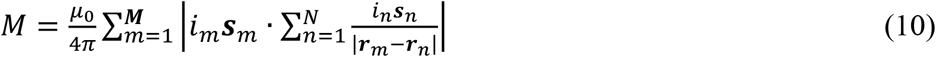

### F. Magnetic Cores

A magnetic material with an isotropic *piecewise constant* relative permeability *μ*_*r*_(***r***) is placed into a primary magnetic field of the coil with the flux ***B***(***r***). Due to magnetization, a secondary field ***H***^*s*^(***r***) appears. The magnetic material can be removed and replaced, in the evaluation of the secondary field ***H***^*s*^(***r***), by the surface bound charge density *ρ*_*s*_(***r***) [27],[28] so that one has

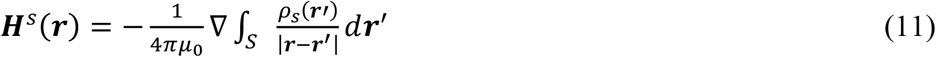

Eq. (11) is identical to the corresponding electrostatic dielectric charge representation (after replacing permeability by permittivity and ***H*** by ***E***) or with the DC current representation (after replacing permeability by conductivity and ***H*** by ***E***). An integral equation for *ρ*_*s*_ – the Fredholm equation of the second kind referred to in [27] as a Phillips-type equation – is obtained using the boundary condition of the continuity of the normal flux across a surface *S* (vasculature vessel boundary) with the local outer normal vector ***n***(***r***) as

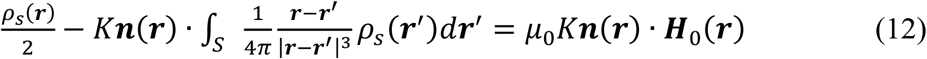

where the magnetic permeability contrast 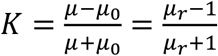 is uniquely defined at the core interface(s). Eq. (12) is solved iteratively and the resulting field from Eq. (11) is added to the primary coil field [28].

## III. Results

### A. Benchmarking Method Speed

Typical computational times of the primary electric field ***E*** and magnetic field ***B*** are reported in [24]. Using a conical-shape coil with 50 single coaxial loops of a circular cross-section and 150,000 elementary current dipoles as an example, the speed of the FMM acceleration was demonstrated. This was accomplished by keeping the coil geometry fixed while varying the number of observation points in a coronal plane whose size is twice the coil size, as shown in Table I and Table II. In these tables, mesh generation time for the wire grid, FMM execution times, and execution times for a plain yet vectorized MATLAB code which directly computes Eq. (2) and Eq. (5) for the observation points are presented. All results have been averaged over several runs. The FMM code indicates only a very modest increase in time for the reported size of the coil model and the sizes of the observation domain. Intel Xeon E5-2698 v4 CPU 2.2 GHz workstations have been used.

**TABLE I.** Timing results for the primary electric field ***E*** computations on 2.2 GHz machines. A coil with 50 single loops, each of which is divided into 100 straight cylinders, and with 30 interpolation points per conductor cross-section is considered.

| Observation Points | Wire Grid Generation Time, s | FMM Execution Time, s | Direct computation time (vectorized MATLAB loop), s |
| --- | --- | --- | --- |
| 100×100 | 0.32 | 0.44 | 92 |
| 200×200 | 0.32 | 0.47 | 394 |
| 300×300 | 0.32 | 0.57 | 892 |
| 500×500 | 0.32 | 0.93 | 2503 |

**TABLE II.** Timing results for the primary electric field ***E*** computations on 2.2 GHz machines. A coil with 50 single loops, each of which is divided into 100 straight cylinders, and with 30 interpolation points per conductor cross-section is considered.

| Observation Points | Wire Grid Generation Time, s | FMM Execution Time, s | Direct computation time (vectorized MATLAB loop), s |
| --- | --- | --- | --- |
| 100×100 | 0.31 | 0.49 | 242 |
| 200×200 | 0.31 | 0.52 | 968 |
| 300×300 | 0.31 | 0.62 | 2250 |
| 500×500 | 0.31 | 0.99 | 6320 |

### B. Example 1. Basic Loop of Current – Comparison Against the Analytical Solution

The first test case was a wire loop of radius 0.05 m and 5 mm cross-sectional diameter, modeled as a set of 3100 elementary current filaments as shown in Fig. 4.

**Fig. 4.**
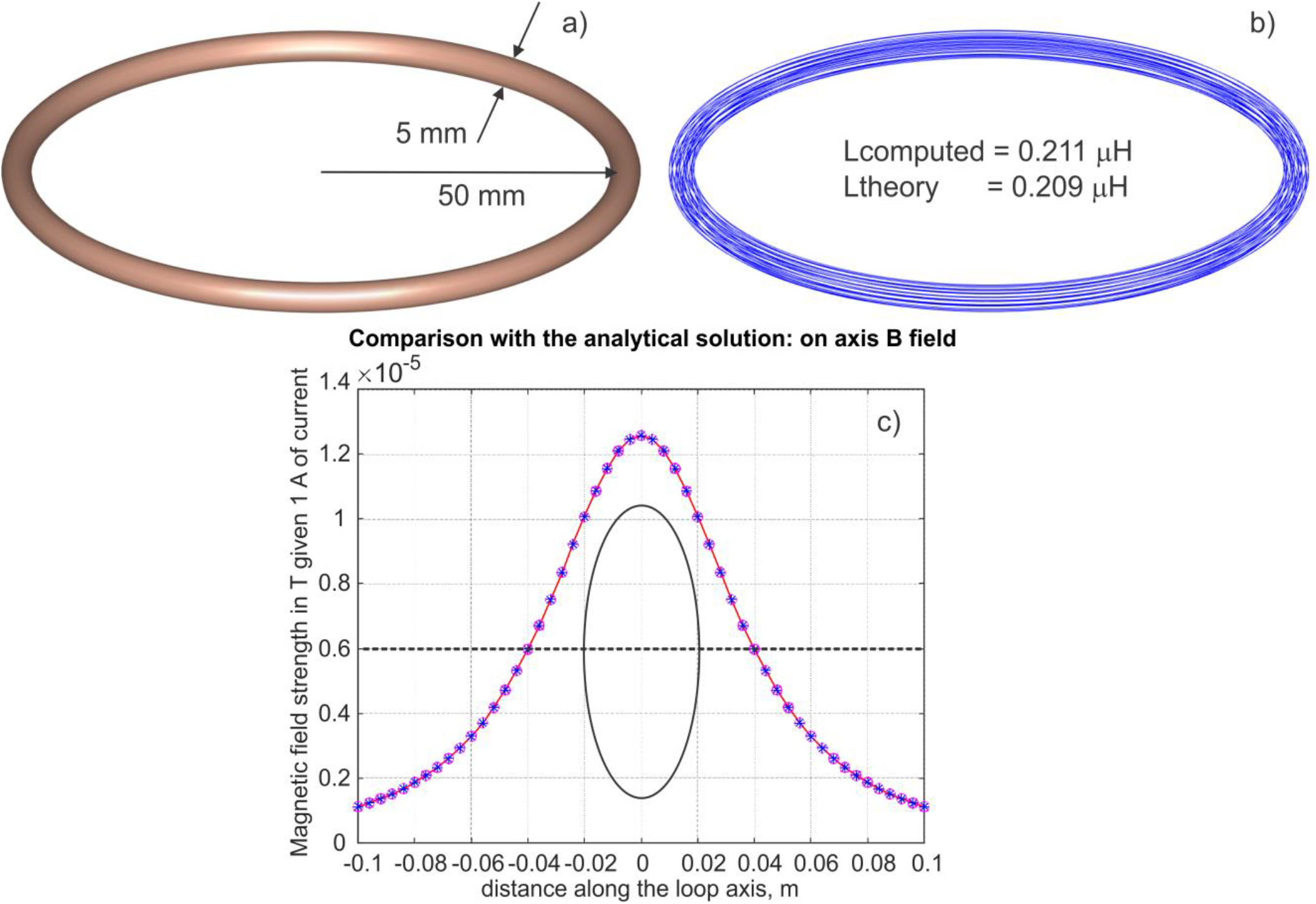
The magnetic field comparison along the axis of a ring of current with the analytical solution and inductance comparison.

The z-component of the magnetic field was computed at 51 observation points on the coil’s axis (−0.1 m < *z* < 0.1 m) via FMM and compared against the analytical solution:

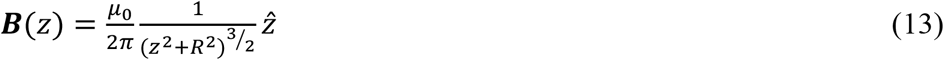

where *R* is the radius of the coil and z is the distance from the plane of the coil to the observation point. For FMM tolerance level iprec = 0, the relative residual error is 2.3 ×10^−4^.

### C. Example 2. BrainsWay H1 Coil – Comparison with Ansys Maxwell 3D FEM Software

To further exemplify the efficiency of the FMM, we modeled the complex geometry of the BrainsWay H1-coil shown in Fig. 5. The coil geometry was created automatically from the known conductor centerline and conductor radius. Fig. 5b displays the 650,000 current filaments that make up the computational coil model. The FMM-accelerated method computed an inductance of 17.503 μH within 1.7 s. Additionally, the electric and magnetic fields were computed over a sphere of radius 0.06 m which was modeled within the coil; the electric field distribution is shown in Fig. 5a.

**Fig. 5.**
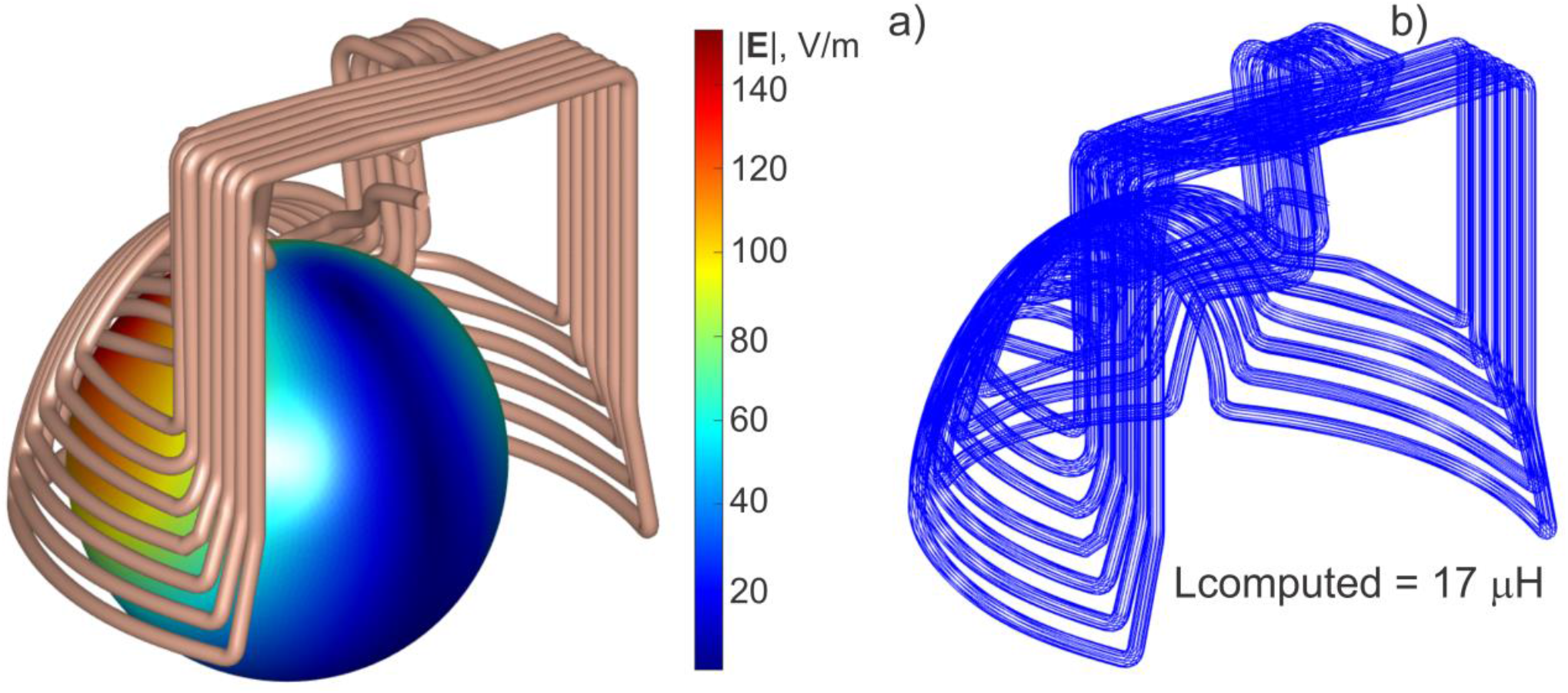
H1-Coil (BrainsWay, US) a) CAD model and electric field over a sphere. b) Computational wire model with only 60,000 of the 650,000 elementary current segments for visibility.

This model was then exported to an STL file and replicated in ANSYS Maxwell 3D (ANSYS Electromagnetic Suite Release 2022 R1) for comparison. The ports were extended to contact the boundary region for appropriate excitation within the Ansys Maxwell interface. The solution ran for 3 hours and 10 minutes (11,420 s) and converged after 8 passes to a relative error of 0.01. The resultant inductance was found to be 17.649 μH - a relative error of 0.83% from the FMM result.

### D. Example 3. Figure Eight Coil with Linear Magnetic Core – Comparison with Ansys Maxwell 3D FEM Software

The third test case was an angled figure-eight coil whose geometry is shown in Fig. 6. The coil was modeled in Ansys and in FMM as a pair of separate loops in series, and the coil’s self-inductance was computed both with and without a magnetic core. The calculated inductances and required solution times for Ansys and FMM are given in Table III. The two methods show excellent agreement in both cases, and the FMM-accelerated method is several hundred to one thousand times faster.

**Fig. 6.**
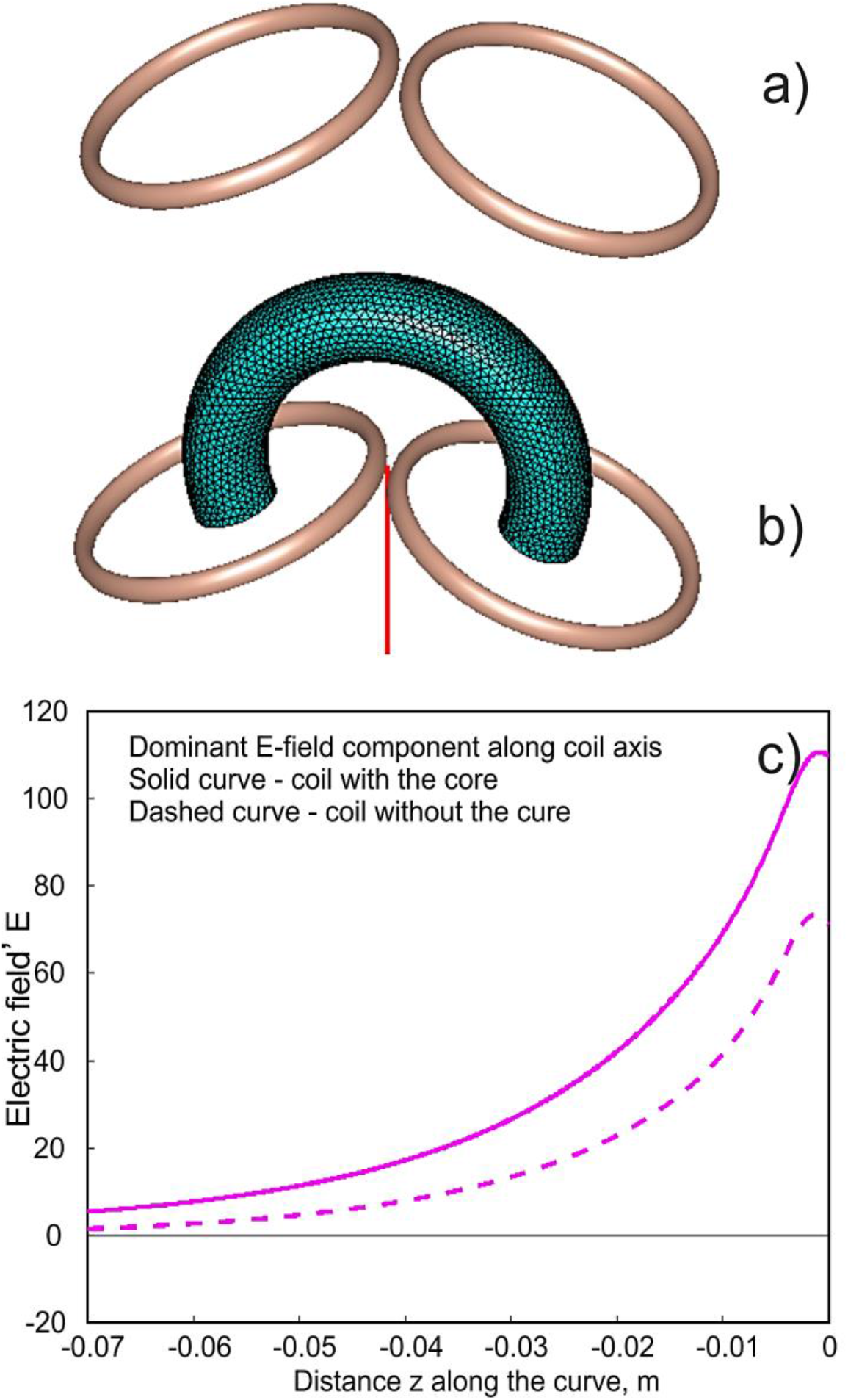
Coil array modeled as two copper coils with an elliptical cross-section. a) CAD model of the coil metal conductor b) Coil with the core, coil axis is shown in red. c) Dominant E-field component on the coil axis assuming linear core.

**TABLE III.**
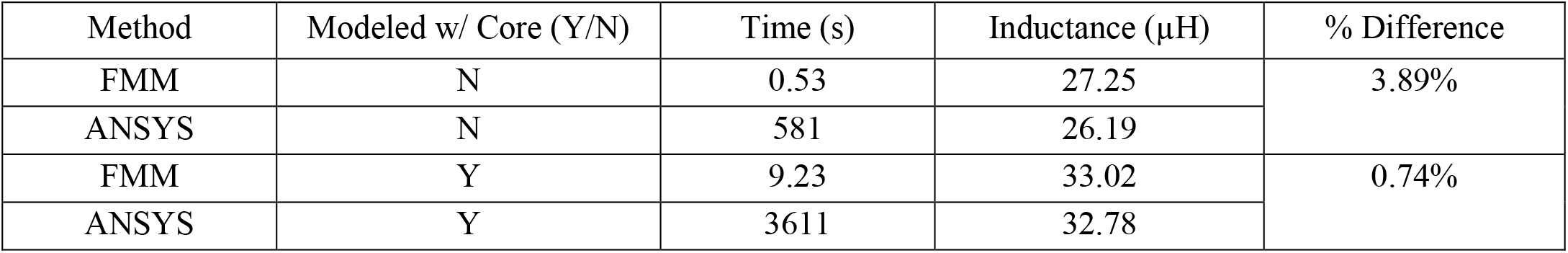
Inductance computations for two separate loops in series, modeled with and without a magnetic core, and compared with Ansys Maxwell 3D.

### E. Example 4. MagVenture C-B60 Coil – Comparison with Measured Values

We validated our model using the MagVenture C-B60 coil with three types of measurements: 1) measurement of coil inductance 2) measurement of physical coil winding dimensions; and 3) measurement of electric field distribution.

First, we measured the inductance of the coil without the coil cable using an LCR meter (NF Corporation ZM2376). At 1 kHz frequency, the measured inductance was 12.76 μH. The inductance of the modeled coil was computed to be 12.60 μH, only a 1.25% difference when compared with the measured inductance.

The coil was dissected with a waterjet cutter (Fig. 7b) in order to obtain exact measurements of the centerline trajectory and cross-section. The winding geometry was digitally replicated, and the same parameters (inductance and E-field) were computed via FMM and compared with the empirically-obtained results. The CAD coil model, computational wire model, and SimNIBS computational coil model for comparison are shown in Fig. 8 on the following page.

**Fig. 7.**
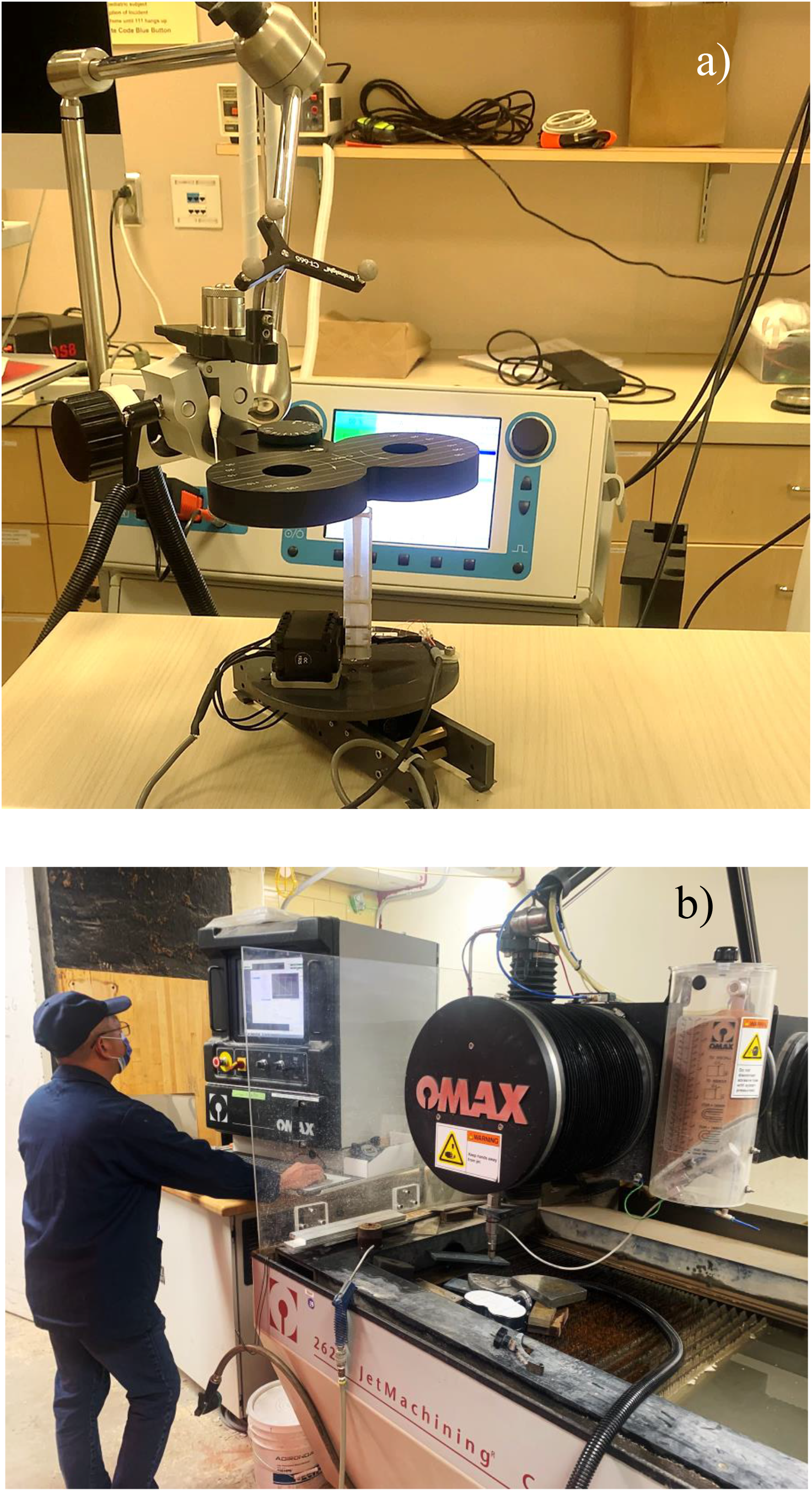
a) NKI E-field triangular probe. b) Waterjet dissection of the MagVenture C-B60 Coil.

**Fig. 8.**
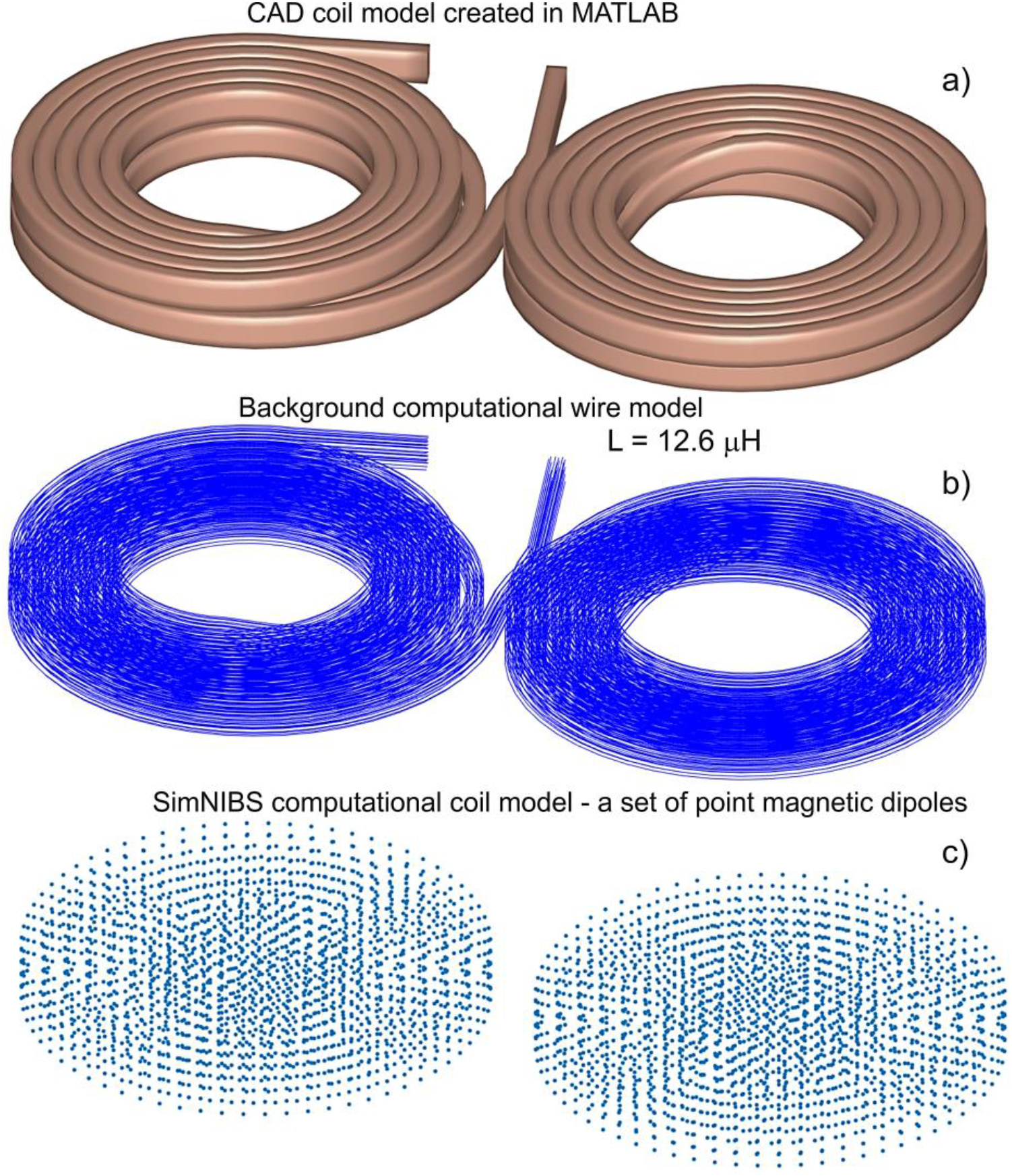
C-B60 coil (MagVenture, Denmark) a) CAD coil model. b) Computational wire model. c) SimNIBS computational coil model represented by a set of point magnetic dipoles.

The electric field was spatially mapped using the NKI triangular E-field probe (Fig. 7a) and procedure as outlined in [18]. The field was sampled at 1000 points on the surface of a 70 mm-radius hemisphere separated from the base of the coil by 20 mm. A MagPro X100 stimulator was used to drive the coil; the rate of change of current was 75e6 A/s, or half of this stimulator’s maximum output.

Next, the FMM was applied to compute the modeled coil’s electric field at the same observation points by exploiting a reciprocal relationship between the NKI probe and a conducting 70 mm radius sphere. Fig. 9 shows the simulated E-field distribution on the measurement surface. Fig. 10 compares the dominant component, *E*_*θ*_, of the measured and simulated E-field at the 1000 observation points. Strong correlation is visible and the 2-norm relative error for the entire hemisphere does not exceed 13%.

**Fig. 9.**
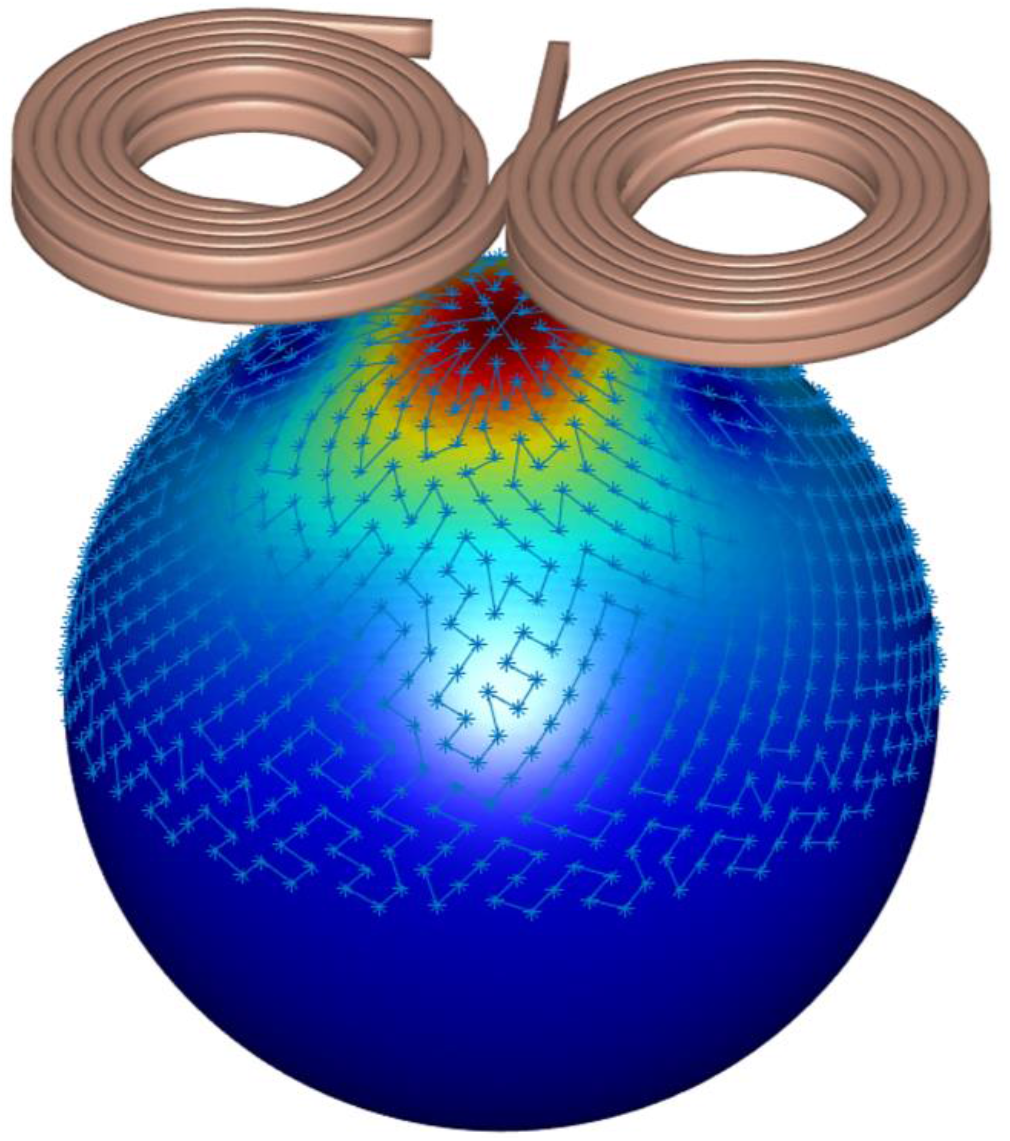
C-B60 coil and electric field over conducting sphere.

**Fig. 10.**
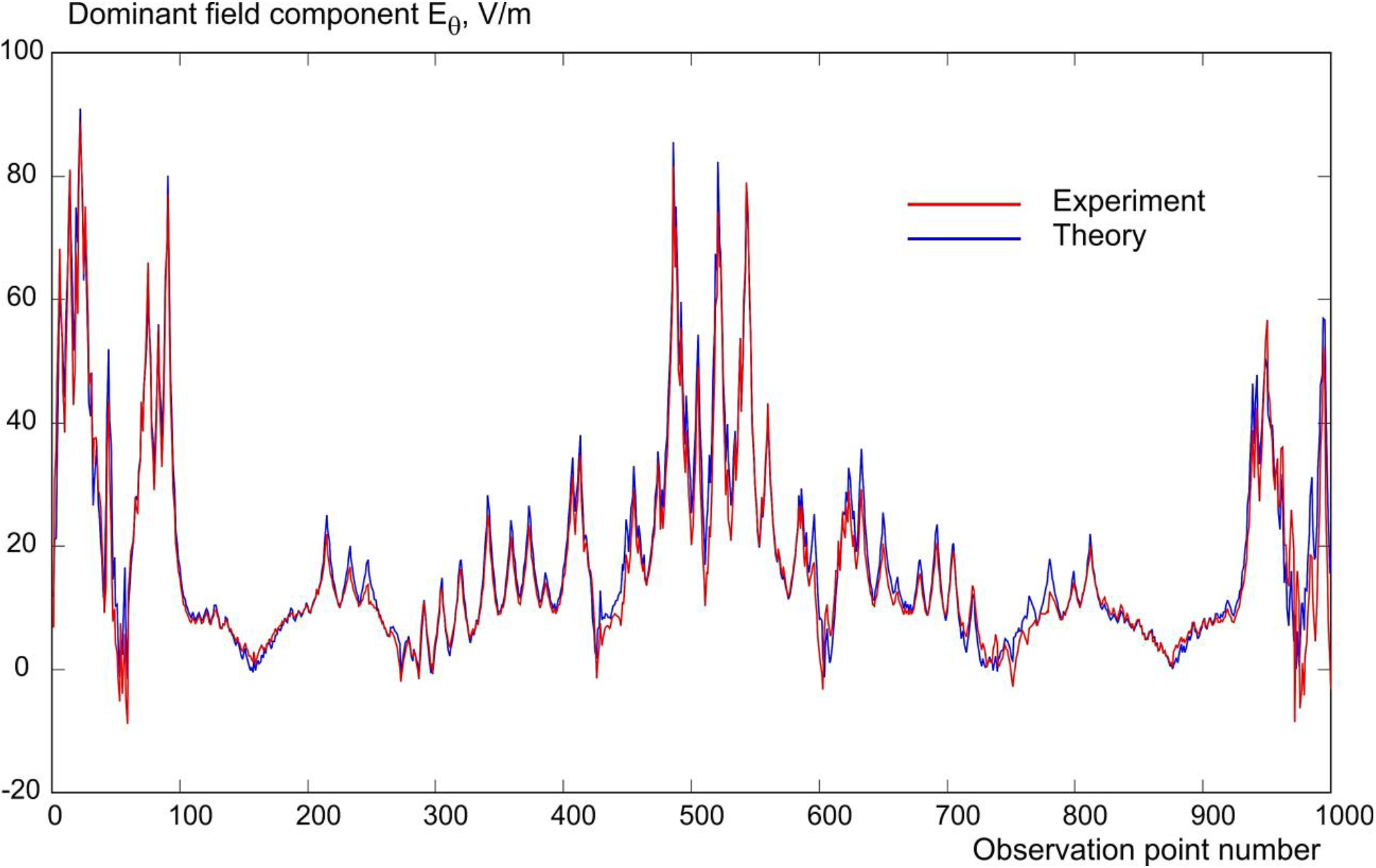
Comparison between the computational and measured dominant field component *E*_θ_ at each observation point.

## IV. Discussion and Conclusions

In this study, an accessible software package for TMS coil analysis and design is described. The user specifies the centerline of the coil conductor as a curve in three-dimensional space, along with the conductor’s cross-section and cross-section type. A computational wire mesh and an illustrative CAD mesh are automatically generated. The wire mesh can model either solid or stranded conductors, with the current flowing either on the surface only or uniformly throughout the conductor’s cross-section, respectively. Using the FMM, electric and magnetic fields of the coils can be determined anywhere in space, and self- and mutual inductances of the coil(s) can be computed.

Recently, Ref. [29] proposed a similar software package, but without mutual inductance calculations, extension to magnetic cores, or fast multipole acceleration – all of which may be useful for several optimization purposes.

The wire-based approach enables precise replication of even the most complex physical conductor geometries, while the FMM acceleration quickly evaluates large quantities of elementary current filaments – both of which are demonstrated in the modeling of the BrainsWay H1 coil in Section III.C. The self-inductance of a model comprising 650,000 current filaments can be computed within 1.7 seconds when using a 2.6 GHz workstation.

This software package also supports modeling of magnetic coil cores, as demonstrated in Section III.D. Agreement to within 0.74% was obtained between the inductances computed by the FMM method and ANSYS Maxwell 3D for the same coil model. Although not provided in this study, it is possible to evaluate non-linear magnetic cores in addition to the linear core exemplified [28].

In addition to comparisons against other numerical methods, an experimental comparison was carried out against a physical MagVenture C-B60 coil as described in Section III.E. The measured and simulated inductances differed by only 1.25%, and nearly perfect correlation was found between the measured and computed E-field values at each observation point.

A limitation of the developed software package is that a uniform current distribution along the distributed filaments is assumed, though future advancements of the package may explore non-uniform current distributions. The program performance scales well with additional CPU cores. Finally, the developed software package can be applicable to any quasistatic inductor design, not necessarily TMS coils only, making it a versatile tool for researchers in a variety of fields. The source code for this software package replicating all examples of the present study can be found online [30].

## Acknowledgements

The authors wish to thank Dr. Samuel Zibman of BrainsWay for providing the H1 coil model. We thank the National Institute of Mental Health (NIMH) Section on Instrumentation for electric field probe construction and coil dissection. This work has been supported by NIH/NIMH grant R01MH130490 and partially supported by NIH/NIMH, NIH//NIBIB, and NIH/NIDCD grants R01MH128421, 5P41EB030006, and 1R01DC020891. Z.-D. Deng is supported by the NIMH Intramural Research Program (ZIAMH002955). H.B. Lu is supported by the NIDA Intramural Research Program (ZIADA000638-01).

## References

[1] Z.-D. Deng, S. H. Lisanby, and A. V. Peterchev, “Electric field depth-focality tradeoff in transcranial magnetic stimulation: simulation comparison of 50 coil designs,” Brain Stimul., vol. 6, no. 1, pp. 1–13, 2013.

[2] Y. S. Cho, H. S. Suh, W. H. Lee, and T.-S. Kim, “TMS modeling toolbox for realistic simulation,” Annu. Int. Conf. IEEE Eng. Med. Biol. Soc., vol. 2010, pp. 3113–3116, 2010.

[3] S. N. Makarov et al., “Preliminary upper estimate of peak currents in transcranial magnetic stimulation at distant locations from a TMS coil,” IEEE Trans. Biomed. Eng., vol. 63, no. 9, pp. 1944–1955, 2016.

[4] P. Rastogi, E. G. Lee, R. L. Hadimani, and D. C. Jiles, “Transcranial Magnetic Stimulation-coil design with improved focality,” AIP Adv., vol. 7, no. 5, p. 056705, 2017.

[5] A. Thielscher and T. Kammer, “Linking physics with physiology in TMS: a sphere field model to determine the cortical stimulation site in TMS,” Neuroimage, vol. 17, no. 3, pp. 1117–1130, 2002.

[6] A. Thielscher, A. Antunes, and G. B. Saturnino, “Field modeling for transcranial magnetic stimulation: A useful tool to understand the physiological effects of TMS?,” Annu. Int. Conf. IEEE Eng. Med. Biol. Soc., vol. 2015, pp. 222–225, 2015.

[7] G. B. Saturnino, O. Puonti, J. D. Nielsen, D. Antonenko, K. H. Madsen, and A. Thielscher, “SimNIBS 2.1: A comprehensive pipeline for individualized electric field modelling for transcranial brain stimulation,” in Brain and Human Body Modeling, Cham: Springer International Publishing, 2019, pp. 3–25.

[8] M. Drakaki, C. Mathiesen, H. R. Siebner, K. Madsen, and A. Thielscher, “Database of 25 validated coil models for electric field simulations for TMS,” Brain Stimul., vol. 15, no. 3, pp. 697–706, 2022.

[9] C. L. W. Sonntag, E. A. Lomonova, J. L. Duarte, “Implementation of the Neumann formula for calculating the mutual inductance between planar PCB inductors,” in 2008 18th International Conference on Electrical Machines, 2008, pp. 1–6.

[10] R. Dengler, “Self inductance of a wire loop as a curve integral,” Adv. Electromagn., vol. 5, no. 1, p. 1, 2016.

[11] S. Liu, J. Su, and J. Lai, “Accurate expressions of mutual inductance and their calculation of Archimedean spiral coils,”Energies, vol. 12, no. 10, p. 2017, 2019.

[12] P. C. Miranda, M. Hallett, and P. J. Basser, “The electric field induced in the brain by magnetic stimulation: a 3-D finite-element analysis of the effect of tissue heterogeneity and anisotropy,”IEEE Trans. Biomed. Eng., vol. 50, no. 9, pp. 1074–1085, 2003.

[13] F. S. Salinas, J. L. Lancaster, and P. T. Fox, “Detailed 3D models of the induced electric field of transcranial magnetic stimulation coils,”Phys. Med. Biol., vol. 52, no. 10, pp. 2879–2892, 2007.

[14] F. S. Salinas, J. L. Lancaster, and P. T. Fox, “3D modeling of the total electric field induced by transcranial magnetic stimulation using the boundary element method,”Phys. Med. Biol., vol. 54, no. 12, pp. 3631–3647, 2009.

[15] I. Laakso, A. Hirata, and Y. Ugawa, “Effects of coil orientation on the electric field induced by TMS over the hand motor area,”Phys. Med. Biol., vol. 59, no. 1, pp. 203–218, 2014.

[16] L. M. Koponen, J. O. Nieminen, and R. J. Ilmoniemi, “Minimum-energy coils for transcranial magnetic stimulation: application to focal stimulation,”Brain Stimul., vol. 8, no. 1, pp. 124–134, 2015.

[17] L. M. Koponen, J. O. Nieminen, T. P. Mutanen, M. Stenroos, and R. J. Ilmoniemi, “Coil optimisation for transcranial magnetic stimulation in realistic head geometry,”Brain Stimul., vol. 10, no. 4, pp. 795–805, 2017.

[18] J. O. Nieminen, L. M. Koponen, and R. J. Ilmoniemi, “Experimental characterization of the electric field distribution induced by TMS devices,”Brain Stimul., vol. 8, no. 3, pp. 582–589, 2015.

[19] P. I. Petrov, S. Mandija, I. E. C. Sommer, C. A. T. van den Berg, and S. F. W. Neggers, “How much detail is needed in modeling a transcranial magnetic stimulation figure-8 coil: Measurements and brain simulations,”PLoS One, vol. 12, no. 6, p. e0178952, 2017.

[20] L. J. Gomez, A. C. Yucel, and E. Michielssen, “The ICVSIE: A general purpose integral equation method for bio-electromagnetic analysis,”IEEE Trans. Biomed. Eng., vol. 65, no. 3, pp. 565–574, 2018.

[21] L. Greengard and V. Rokhlin, “A new version of the Fast Multipole Method for the Laplace equation in three dimensions,” cta Numer., vol. 6, pp. 229–269, 1997.

[22] “Fast multipole methods in three dimensions (FMM3D) — fmm3d 1.0.0 documentation,”Readthedocs.io. [Online]. Available: https://fmm3d.readthedocs.io/en/latest/. [Accessed: 11-May-2023].

[23] B. K. Choi and C. S. Lee, “Sweep surfaces modelling via coordinate transformation and blending,”Comput. Aided Des., vol. 22, no. 2, pp. 87–96, 1990.

[24] S. N. Makarov, L. N. de Lara, G. M. Noetscher, and A. Nummenmaa, “Modeling primary fields of TMS coils with the fast multipole method,” 2019.

[25] C. A. Balanis, Advanced Engineering Electromagnetics, 2nd ed. Chichester, England: John Wiley & Sons, 2012.

[26] J. D. Jackson, “Classical Electrodynamics,” 3rd ed., Nashville, TN: John Wiley & Sons, 1975, p. 261.

[27] J. G. Van Bladel, Electromagnetic Fields. John Wiley & Sons, 2007.

[28] S. N. Makaroff, H. Nguyen, Q. Meng, H. Lu, A. R. Nummenmaa, and Z.-D. Deng, “Modeling transcranial magnetic stimulation coil with magnetic cores,”J. Neural Eng., vol. 20, no. 1, p. 016028, 2023.

[29] M. Köhler and S. Götz, “TMS coil design instrument (Kl/Codein Box): A toolbox for creating user-defined coils from conductor path data,”Brain Stimul., vol. 16, no. 3, pp. 698–700, 2023.

[30] L. Morales et al., “Compiled Coilo Code,” Dropbox. [Online]. Available: https://www.dropbox.com/sh/ta2vg50ze9rp6c9/AAD_kWjcarg3icVK6QSvfn65a?dl=0.

